# Ethylene Glycol Monomethyl Ether Altered Rat Sperm Small RNAs with Critical Developmental Roles

**DOI:** 10.64898/2025.12.29.693789

**Authors:** Yuan Pu, August Guang, Xinran Qi, Poonam Mehta, Saadhya Bahudodda, Angela R. Stermer, Daniel J. Spade

**Affiliations:** Center for Computational Biology of Human Disease, Brown University, Providence, RI, USA; Department of Pathology and Laboratory Medicine, Brown University, Providence, RI, USA

**Author notes:** To whom correspondence should be addressed at: Brown University, Department of Pathology and Laboratory Medicine, Box G-E5, Providence, RI, USA. Authors contributed equally to the manuscript (Yuan Pu, August Guang).

**Keywords:** testis, germ cells, spermatozoa, RNA, sncRNA

## Abstract

Ethylene glycol monomethyl ether (EGME) is a testicular germ cell toxicant that selectively targets spermatocytes. In rats, male-only EGME exposure reduces mating success and can lead to an increase in resorbed fetuses. In a previous study, five-day exposure to 50, 60, or 75 mg/kg/d EGME in male rats led to a decrease in sperm motility and increase in retained spermatid heads with a lowest observed adverse effect level (LOAEL) of 75 mg/kg/d. At 60 mg/kg/d, EGME exposure altered the proportion of sperm small RNA reads mapped to different small RNA categories and the distribution of read lengths. To understand the possible role of sperm sncRNAs in EGME-mediated reduction in spermatogenesis, fertility, and embryonic development, we analyzed sperm sncRNA data from EGME-treated male rats to identify differential expression at the individual RNA level. EGME treatment resulted in dose-dependent increases in the expression levels of microRNAs (miRNAs), piRNAs, and tRNA-derived small RNAs (tsRNAs), and mixture of dose-dependent increases and decreases in abundance of rsRNA reads. We identified 12 miRNAs that were differentially expressed at all EGME doses, with a monotonic, dose-dependent increase. High-confidence targets of these 12 miRNAs are known to be expressed in pre-implantation embryos and statistically enriched for Gene Ontology (GO) biological processes related to early development, such as cell fate commitment and regulation of developmental growth. These results demonstrated that the EGME-induced changes in sperm sncRNA levels were reproducible, dose-dependent, and provided a putative mechanism of paternal EGME effects on embryonic development, which will be investigated in future studies.

## Introduction

Ethylene glycol monomethyl ether (EGME) is glycol ether used as a plasticizer, deicer, and in inks and dyes (Dieter 1993), and is a well-characterized testicular toxicant. Primary exposure concerns are occupational, and it is regulated in the United States by OSHA (OSHA) and the U.S. EPA (U.S. EPA 2022). Upon oral exposure, EGME is metabolized to the active metabolite, 2-methoxyacetic acid (MAA) (Beattie and Brabec 1986). EGME exposure results in mitochondrial dysfunction (Foster et al. 1983) and subsequent apoptosis of germ cells, with preference for pachytene and meiotic spermatocytes (Creasy and Foster 1984; Creasy et al. 1986). EGME-induced germ cell death is rapid, detectable at 12 hours after a dose of 250 mg EGME/kg body weight (Creasy et al. 1986) and 24 hours after a dose of 150 mg EGME/kg body weight in rats (Chapin et al. 1984). Perhaps more interestingly, EGME is an example of a chemical that is not directly genotoxic, i.e. is negative in the Ames assay (Hoflack et al. 1995) and other tests for DNA damage (McGregor 1984), but for which paternal exposure causes adverse effects on embryo development. Short-duration exposure to EGME in rats leads to post-implantation embryo loss that is delayed by five weeks (Chapin, Dutton, Ross, and Lamb 1985), a time frame that corresponds with the time it takes for pachytene spermatocytes to mature into spermatozoa and complete epididymal maturation.

The mechanisms of paternal preconception exposure-induced developmental toxicity are not well understood. Because there is evidence that EGME and its major metabolite, MAA, are not genotoxic, it follows that EGME has the potential to disrupt embryo development through an epigenetic mechanism. One plausible epigenetic mechanism is disruption of sperm RNA. Sperm contains mRNA, and small non-coding RNAs (sncRNA), including miRNA, piRNA, and tsRNA, which have putative roles in embryo development (Sendler et al. 2013; Jodar et al. 2015; Conine et al. 2018). These sperm RNAs are well-conserved across mammalian models; human, mouse, and rat sperm all contain thousands of mRNAs, with one report identifying 6,684 transcripts expressed in all three species (Bianchi et al. 2021). In particular, there is evidence for a critical role of miRNAs and endo-siRNAs in the male germline. Germ-cell specific loss of *Dicer* or *Dgcr8* from embryonic day 15, driven by Ddx4-Cre in mice, leads to infertility and severe reduction of meiotic progression and maturation of elongate spermatids (Zimmermann et al. 2014). Furthermore, there is increasing evidence that altered levels of regulatory RNAs in sperm contribute to disease phenotypes such as metabolic disorder in offspring (Zhang et al. 2019).

EGME alters testicular gene expression and miRNA expression, which suggests that sperm RNAs may be altered following EGME exposure, as well. In monkeys exposed to EGME, expression of many testicular miRNAs increases, while serum miRNA levels decrease (Sakurai et al. 2015). In rats, EGME exposure results in changes in expression of meiosis-related genes in the testis (Tonkin et al. 2009). Further, we previously reported that low-dose EGME alters the proportions and read length distributions of several classes of small RNA in rat sperm (Stermer et al. 2019). Rats were treated with 50-75 mg EGME/kg body weight/d for 5 days. After 5 weeks with no treatment, no testicular histopathology was observed at doses lower than 75 mg/kg/d.

However, at 50 mg/kg/d, the proportion of miRNA reads relative to all small RNA reads increased, and the distribution of piRNA and tsRNA read length was significantly altered. Here, we conducted a new analysis of the Stermer et al. (2019) dataset to determine the impact of EGME exposure on expression of specific sperm small RNAs, with the goal of identifying dose-dependent regulatory RNA changes in sperm. These sensitive, dose-dependent changes have the potential to serve as biomarkers of exposure to EGME and possibly other toxicants, as well as candidate RNAs in the mechanisms of developmental toxicity of paternal preconception EGME exposure.

## Materials and Methods

### sncRNA-seq Data

Raw small RNA sequencing data were obtained from a previously published study (Stermer et al. 2019). Briefly, Stermer et al. (2019) exposed adult male Fisher rats to EGME according to an established protocol (Chapin, Dutton, Ross, and Lamb 1985). Rats were exposed to 0, 50, 60, or 75 mg/kg EGME/d by oral gavage, in water vehicle, for five days. After the five-day treatment period, no further treatment was administered for five weeks. This is the approximate time required for pachytene spermatocytes, which are the target of EGME, to develop into mature spermatozoa, and has been established as the time when the magnitude of EGME reproductive toxicity is greatest (Chapin, Dutton, Ross, and Lamb 1985). Prior experiments established that the lowest observed adverse effect level (LOAEL) for fertility effects of EGME in this experimental paradigm was 100 mg/kg/d (Chapin, Dutton, Ross, and Lamb 1985), while the LOAEL for reduced sperm motility was 75 mg/kg/d (Stermer et al. 2019). After five weeks, rats were euthanized, and spermatozoa were obtained from the cauda epididymis by repeated puncture and washing. Sperm small RNA was prepared using an established protocol in which sperm samples are subjected to somatic cell lysis and washing prior to isolation of RNA using the miRvana kit (ThermoFisher Scientific). This protocol is well-validated and results in total spermatozoa RNA free of rRNA from somatic cells or early germ cells, based on Agilent 2100 Bioanalyzer analysis, and free of markers of immune and epithelial cells, *CD45* and *CDH1*, respectively, while expressing the RNA coding for the sperm protamine protein, *PRM1* (Bianchi et al. 2018). We have previously reported using this method to obtain RNA from rat, mouse, and human samples (Bianchi et al. 2019; Bianchi et al. 2021; Qi et al. 2025). Subsequently, RNA sequencing libraries were prepared from 10 ng small RNA/sample using the Ion total RNA seq kit v2 (ThermoFisher Scientific), and single-end sequencing of small RNA was performed on the 39 libraries (one sample in the 60mg/kg exposure group lacked sufficient RNA to process) with an IonProton sequencer (ThermoFisher Scientific). This yielded an average of 16 million reads per library and an average read length of 30 nt. Raw sequence reads are available from the NIH Sequence Read Archive database (accession no. PRJNA492909).

### Small RNA Expression Profiling

Initial quality control was performed using FASTQC (Andrews 2010). Two samples with extremely low read counts were identified as outliers and excluded from downstream analysis. Small RNA expression levels were then profiled using sRNAbench (Aparicio-Puerta et al. 2022) in genome mode. Reads were mapped against the rat genome (Rnor_6_0) and compared to miRBase Release 22.1 (Kozomara et al. 2019). Default sRNAbench parameters were applied: the bowtie seed alignment, seed length 20, minimum read count 2, minimum read length 15, allowed 1 mismatch, and 10 multiple mappings. After expression profiling, multi-map adjusted read counts for small RNAs (miRNAs, tRNAs, ncRNAs, cDNA) from sRNAbench “.grouped” files were parsed using a custom Python script.

### Differential Expression Analysis

Differential expression of small RNAs across dosage groups was determined using DESeq2 (Love et al. 2014) with default filtering. A given small RNA passed the filter if it had at least 9 samples with >10 counts each for the RNA. Both the Wald test and the likelihood ratio test were run for each small RNA type. For miRNAs, significantly differentially expressed miRNAs (padj < 0.05) were identified for each of the three dosage vs. control pairs (50mg/kg vs 0 mg/kg, 60mg/kg vs 0 mg/kg, and 75mg/kg vs 0mg/kg). The intersection of these significant miRNA sets across all comparisons was used for further analysis. The raw and normalized count data are available from the NIH Gene Expression Omnibus database (accession no. GSE313131). Small RNA reads mapped to the RNACentral rRNA annotations in sRNAbench were assigned to 8 unique rRNAs and analyzed for differential abundance using FUSION (Rawal et al. 2025).

As RNAseq methods such as DESeq2 have high false positive rates (Li et al. 2022), we assessed robustness of our results with a Monte Carlo simulation (Dere et al. 2016; Dere et al. 2018). We created 1000 subsamples of our dataset where each subsample consisted of smaller groups of 6 individuals in each group, where the individuals were sampled without replacement from the larger set of 10 individuals per group. We then ran DESeq2 again on each of the 1000 subsamples and counted how many times we found the same differentially expressed miRNAs. *Target Gene Prediction and Functional Enrichment*

Putative target genes for the intersection set of significant miRNAs were assembled by manually querying each miRNA in miRDB (Chen and Wang 2020). Targets with scores greater than 80 were retained, yielding 2073 unique genes. An over-representation analysis (ORA) was performed on the full list of target genes, using the R package ClusterProfiler (Yu 2024). The default background gene list (whole rat genome) was used, as gene expression was not measured in this experiment. All three subontologies - biological process (BP), molecular function (MF), and cellular component (CC) were investigated. All the other parameters are left as default. For each subontology, we obtained a list of GO terms ranked by p-values.

## Results

### EGME dose-dependently increased sperm miRNA levels

A total of 405 miRNAs were initially identified by sRNAbench, with 277 remaining after filtering in DESeq2. Sperm miRNA levels showed a high degree of clustering by treatment level, based on Euclidean distance (Fig. 1A), with two main clusters forming, consisting of (1) all 10 vehicle controls, four 50 mg/kg/d EGME samples, and two 60 mg/kg/d EGME samples; and (2) the remaining six 50 mg/kg/d samples, seven 60 mg/kg/d samples, and all ten 75 mg/kg/d samples. In principal component analysis (PCA) (Fig. 1B), there was a clear gradient from vehicle to 75 mg/kg/d along the first principal component. The first two principal components explained 25 and 13% of the variance in the data, respectively, and the samples became more variable within treatment group as the treatment level increased, suggesting that increased EGME-induced germ cell stress led to greater variability. There was distinct visual separation in the PCA plot between sample groups, in particular the vehicle control and 50 mg EGME/kg/d, with less separation between the 60 and 75 mg EGME/kg/d groups.

**Figure 1.**
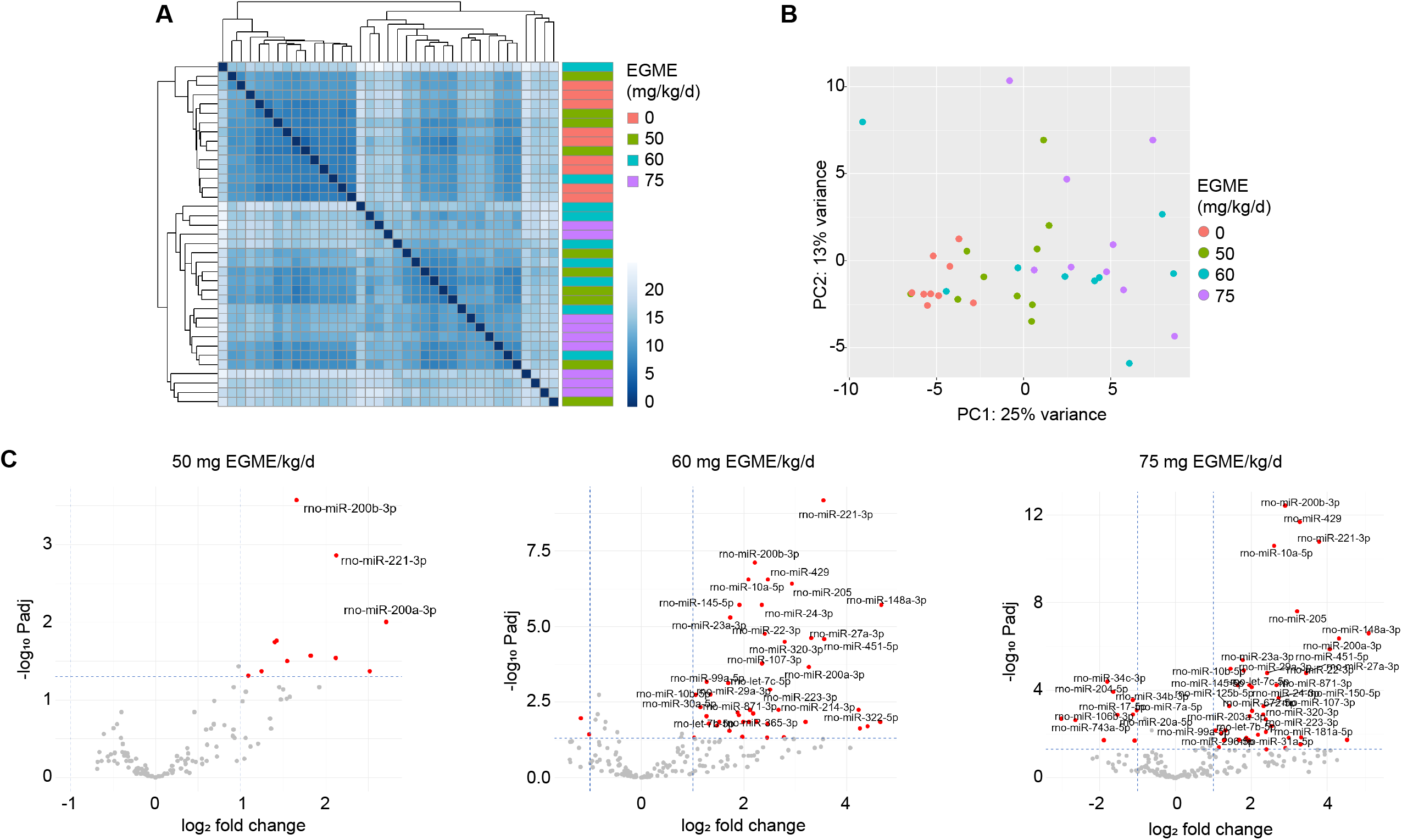
Changes in sperm miRNA levels following EGME exposure. **A**. Hierarchical clustering of EGME- and vehicle control-treated samples showing Euclidean distance based on sperm miRNA read counts. **B**. Principal component analysis of EGME- and vehicle-control-treated samples based on sperm miRNA read counts. **C**. Differential expression of sperm miRNA between vehicle control and 50, 60, or 75 mg/kg/d EGME-treated samples (DESeq2 with the Wald test, p-adj < 0.05, fold change > 2).

The total number of DE miRNAs increased from 12 in the 50 mg/kg/d group to 56 in the 60 mg/kg/d group and 57 in the 75 mg/kg/d group, with the majority of significant DE miRNAs being upregulated in the EGME treatment groups (Fig. 1C, Table S1). There was significant overlap between the treatment groups, with 12 miRNAs being shared among all three groups, and a further 25 DE miRNAs shared by the 60 and 75 mg/kg/d groups (Fig. 2A). A Monte Carlo analysis found that these 12 shared miRNAs were highly stable among the DE miRNAs (Fig. 2B). In 1000 simulations, 10 of the 12 shared DE miRNAs were in the top 12 most frequently identified DE miRNAs, appearing in 748 to 992 out of 1000 iterations. The remaining two shared DE miRNAs were also in the top 40 most stable DE miRNAs. The 12 shared DE miRNAs were miR-200b-3p, miR-221-3p, miR-200a-3p, let-7c-5p, miR-429, miR-365-ep, miR-27a-3p, miR-205, miR-148-3p, miR-23a-3p, miR-24-3p, and let-7b-5p. All 12 monotonically increased across the dose range (Fig. 2C). These 12 miRNAs include members of the Mir-23-27-24 cluster and the Mir-200 family, which are known to be involved in embryonic development (Korpal and Kang 2008; Sundararajan et al. 2022; Yap and Chen 2024). As an additional test of the dose-dependency of these DE miRNAs, we performed the likelihood ratio test in DEseq2, which tests for a significant trend across all treatment groups. There were 56 DE miRNAs by the likelihood ratio test (padj<0.05), which included all 12 core, dose-dependent DE miRNAs from the Wald test (Table S2). To determine whether the miRNAs detected in our experiment were consistent with other reports, we compared our miRNA list with supplemental data from two recent rat sperm RNA experiments (Suvorov et al. 2020; McSwiggin et al. 2024). Of the 277 sperm miRNAs that we report, 190 were reported as DE after exposure to at least one environmental toxicant: atrazine, DDT, jet fuel, or vinclozolin (McSwiggin et al. 2024), and 270 were reported as detected in rat sperm by Suvorov et al. (Suvorov et al. 2020) (Table S3). By comparing the baseMean ranks within each dataset and our own, we found that our data correlated significantly with the Suvorov et al. dataset (Sperman r = 0.5746, p<0.0001), but not the McSwiggin et al. dataset (r = 0.06038, p=0.4080). This may be because of experimental differences such as the use of archival samples in the McSwiggin et al. experiment, or reporting differences, as the McSwiggin et al. paper did not report baseMean values for miRNAs that were not DE. Overall, this analysis confirmed that the sperm miRNAs detected in our experiment were largely consistent with recent literature.

**Figure 2.**
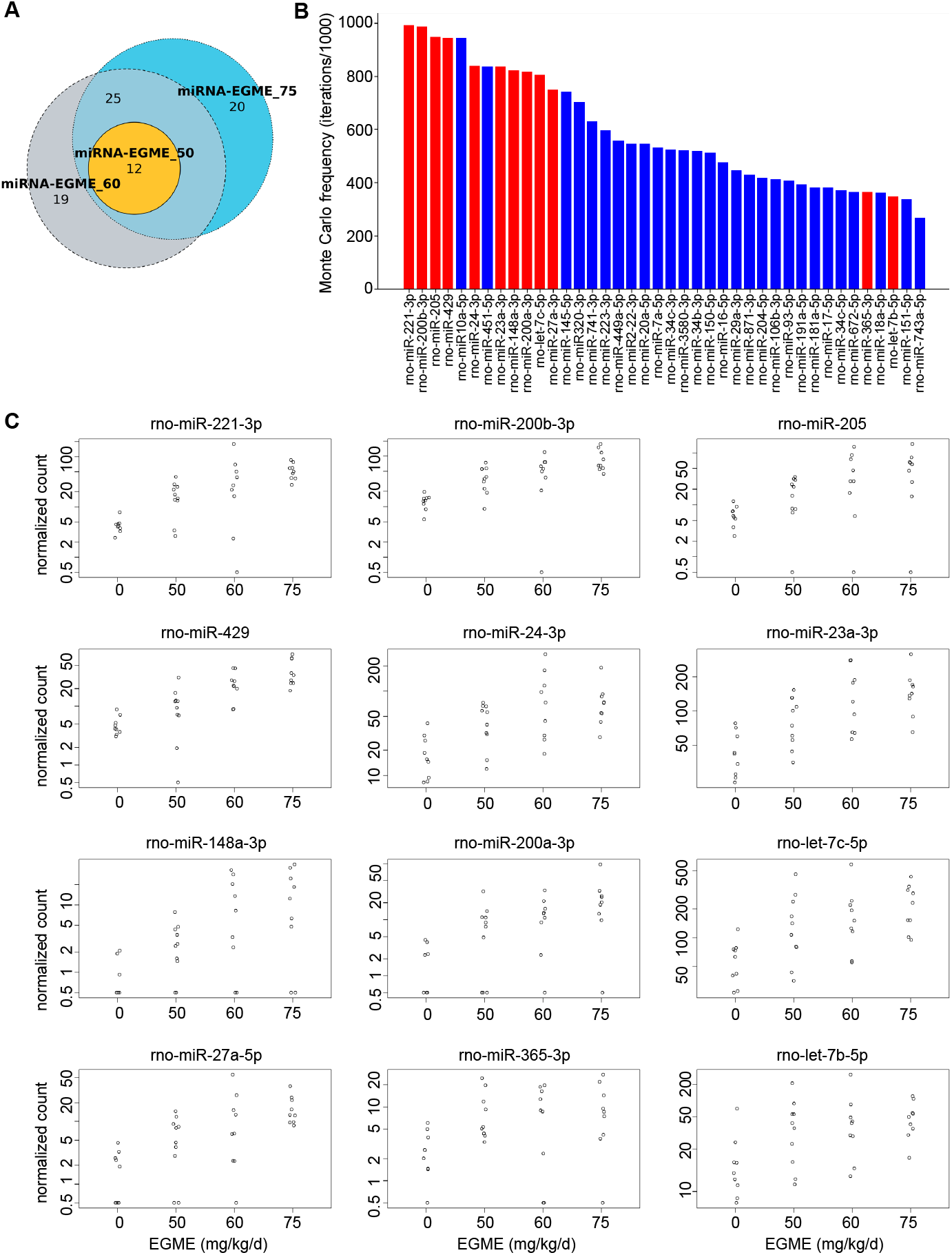
EGME dose-dependency of sperm miRNAs. **A**. Venn diagram showing overlap of differentially expressed miRNAs in comparisons between control and 50, 60, and 75 mg/kg/d EGME. 12 miRNAs were common to all three treatment levels. **B**. Top 40 most commonly differentially expressed sperm miRNAs in a Monte Carlo analysis. Ten of the 12 common differentially expressed miRNAs (shown in red) were in the top 12, and the remaining two were within the top 40 most reproducibly differentially expressed miRNAs. **C**. Normalized read counts for the 12 common differentially expressed miRNAs. In all cases, the response trend was monotonic increase with increasing dose.

### EGME dose-dependent miRNA targets in embryo development

We used miRDB to identify the high-confidence targets of the 12 EGME dose-dependent DE miRNAs, using a miRDB target score > 80 as the cutoff. At this level of confidence, we identified 1,203 targets (Table S4). 787 of the 1,203 target genes were only targets of one miRNA. A further 343 were targets of two miRNAs, and 50, 12, and 7 genes were identified as shared targets of 3, 4, or 5 of the 12 core miRNAs, respectively (Fig. 3A). The most highly shared target genes were *Zeb1, Mmd, Clasp2, Hectd2, Wdr37, Sema6d*, and *Slc6a1*, which were each identified as a target of 5 of the 12 shared DE miRNAs. To assess whether these target genes are likely to have roles in embryonic development, we compared the targets with mRNAs enriched in the epiblast (EPI), primitive endoderm (PE), and trophectoderm (TE) of human e5-7 embryos (Petropoulos et al. 2016). The targets matched 54, 31, and 36 of the EPI, PE, and TE-enriched genes, respectively (Fig. 3B, Table S4), which confirms that the EGME dose-dependent miRNAs target genes involved in early development and that at least a subset of those genes are expressed in early embryos. That subset included well-known early embryo development genes, such as *Aldh1a1, Foxa2, Gata2, Gata4, Gata6*, as well as fundamental genes involved in cell cycle control, cytoskeleton structure and remodeling, and cell signaling. Using the 1,203 target transcripts as input, we identified 321 enriched Gene Ontology (GO) terms (Table S5), including 295 biological process (BP), 9 cellular compartment (CC), and 17 molecular function (MF) terms (Fig. 3C-D). The top 20 GO BPs included important developmental functions, such as “axonogenesis,” “regulation of developmental growth,” “cell fate commitment,” “morphogenesis of a branching structure,” and “morphogenesis of a branching epithelium” (Fig. 3D). Genes under the “cell fate commitment” term, as an example, have well-characterized roles in development. These include *Cyp26b1, Dmrt3, Foxa2, Gata2, Gata4, Gata6, Hoxd10, Kdr, Pax6, Pou3f2, Pparg, Runx1, Runx2, Smad4, Tgfb2, Tgfbr1, Wnt9a*, and *Wnt9b*, among 40 target genes (Table S5). Analysis of target gene sharing among the 12 miRNAs in the GO BP gene sets revealed a high degree of overlap. Of the significant GO BPs, the majority included targets of 6 or more miRNAs, with only one GO BP uniquely linked to a single miRNA (Fig. 3C). 90 out of 295 pathways were shared by all 12 miRNAs, although most target genes were not shared by a large number of miRNAs (Fig. 3A). Beyond biological processes, the top GO CCs included compartments associated with cell division, including “cleavage furrow,” and “cell division site,” as well as terms related to phosphatases, Golgi, and neuronal synapses (Fig. 4A). The top GO MFs included several terms related to kinase and phosphatase activity, transcription, RNA binding, and SMAD binding (Fig. 4B).

**Figure 3.**
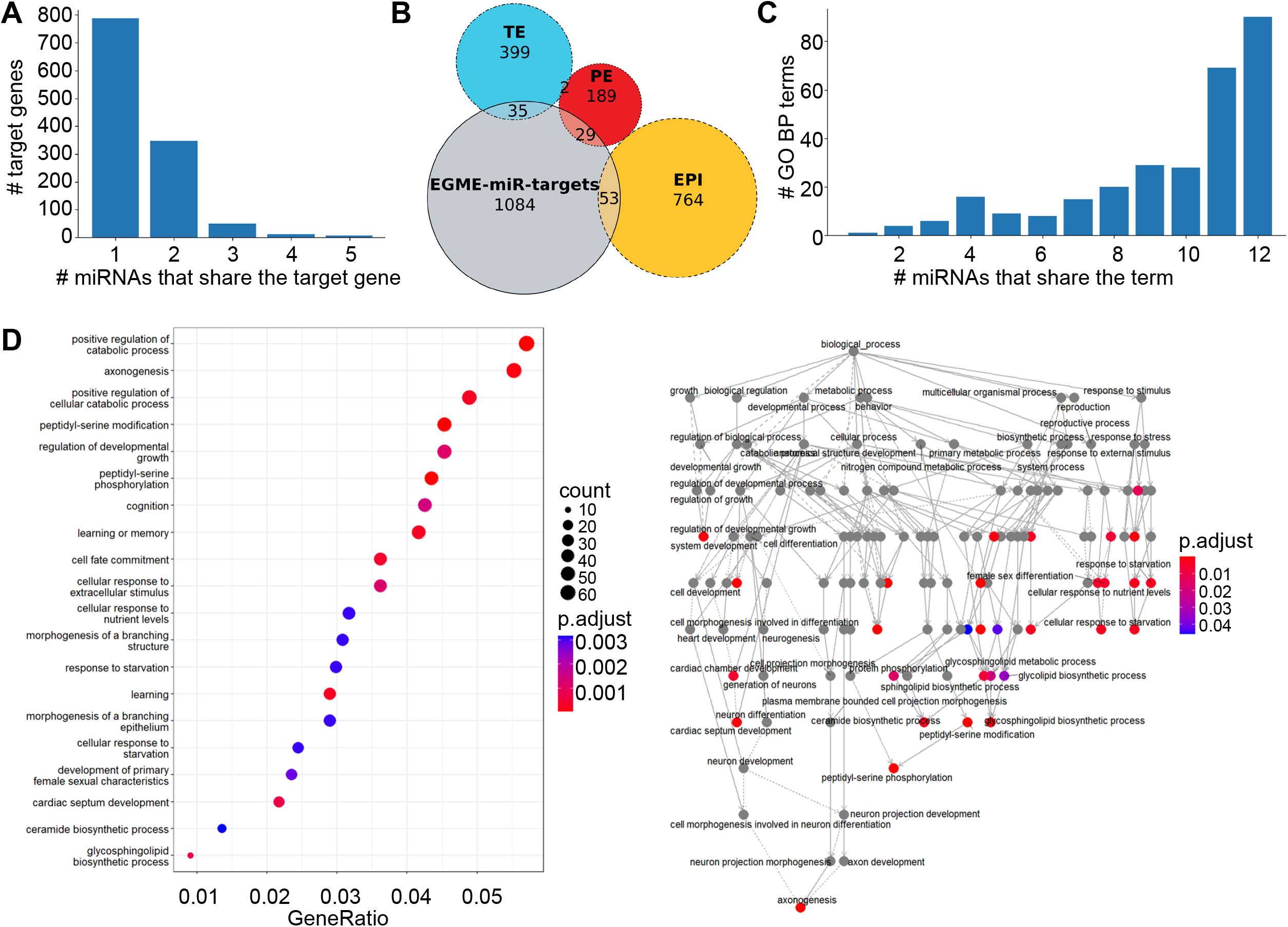
Targets of EGME dose-dependent miRNAs. Using mirDB, we identified 1,203 high-confidence targets of the 12 common differentially expressed miRNAs. **A**. Histogram showing the distribution of target genes by the number of DE miRNAs that share the target. **B**. Venn diagram showing the overlap of the 1,203 common DE miRNA target genes with genes known to be expressed in the epiblast (EPI), primitive endoderm (PE), or trophectoderm (TE) of the early human embryo, from a single-cell RNA-seq analysis (Petropoulos et al. 2016). The analysis identified a total of 119 early human embryo genes among the 1,203 targets of common EGME differentially expressed miRNAs. Note that one gene shared by the EGME-miR-target, PE, and EPI gene list and one shared by the EGME-miR-target, PE, and TE gene lists is not accounted for in the diagram because of graphing constraints. **C**. Histogram showing the number of core miRNAs with target genes common to each of the enriched GO BPs. **D**. Enrichment analysis of Gene Ontology Biological Process terms (GO BPs) for the 1,203 targets of the 12 common differentially expressed miRNAs. Right panel shows hierarchical arrangement of enriched GO BPs.

**Figure 4.**
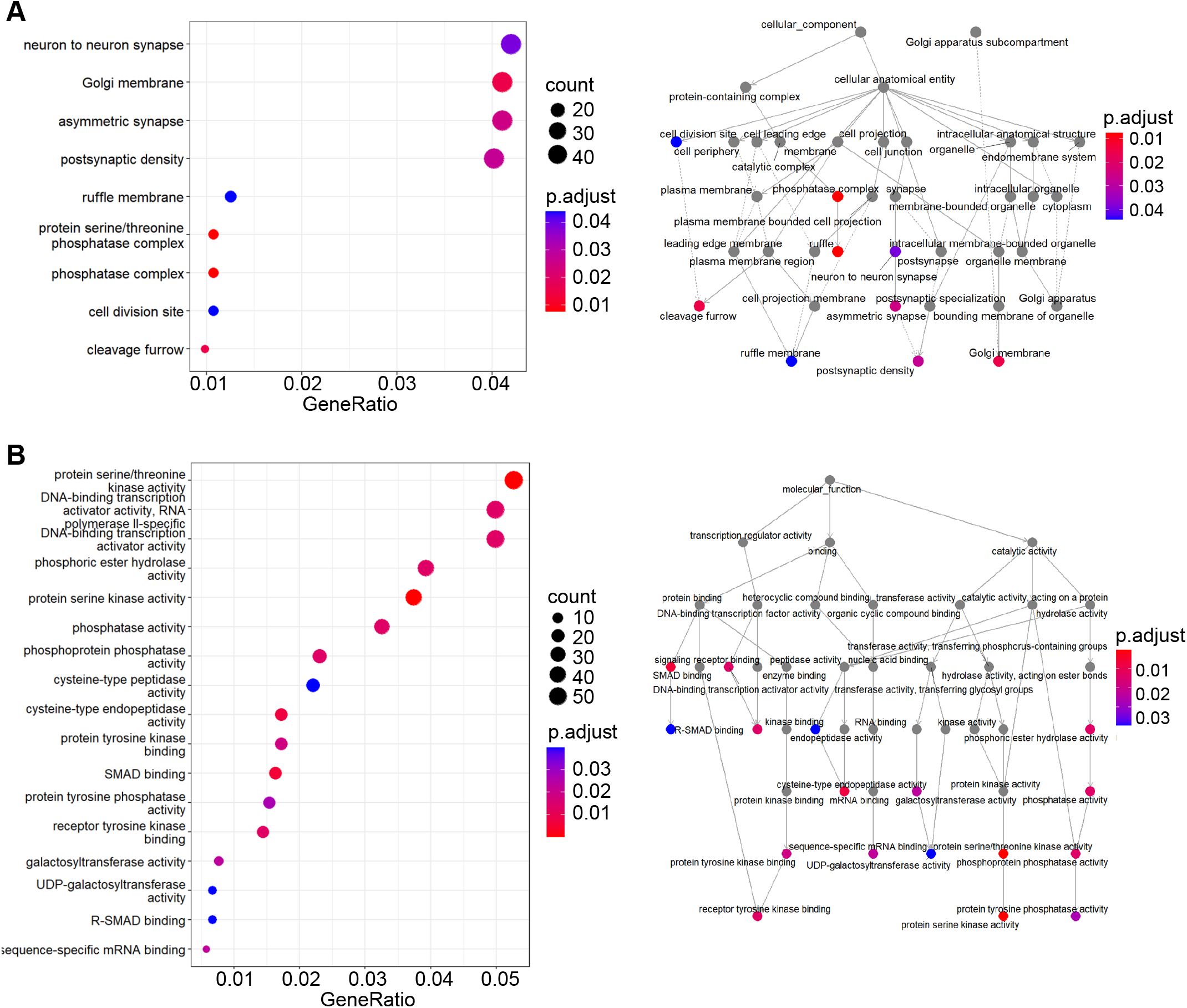
Functional analysis of EGME dose-dependent miRNA targets. Using the 1,203 high-confidence targets of the 12 common differentially expressed miRNAs. **A**. Enrichment analysis of Gene Ontology Cellular Component terms (GO CCs) for the 1,203 targets of the 12 common differentially expressed miRNAs. **B**. Enrichment analysis of Gene Ontology Biological Molecular Function terms (GO MFs) for the 1,203 targets of the 12 common differentially expressed miRNAs. Right panels show hierarchical arrangement of enriched GO MFs.

### EGME effects on sperm tsRNA levels

Small RNA reads mapped to 365 tRNA genes in sRNAbench, which corresponded with 41 unique tRNAs (Fig. 5, Table S6), after removing duplicate tRNA genes (Iben and Maraia 2014). We performed unsupervised analyses of tRNA reads, which showed separation of samples by tRNA expression, similar to the pattern observed with miRNAs. In hierarchical clustering based on Euclidean distance (Fig. 5A), there were four main clusters, showing a lot of separation between the control and high dose group. One cluster contained eight of nine 75 mg/kg/d samples and three of nine 60 mg/kg/d samples, while another contained seven of nine vehicle control samples, four 50 mg/kg/d samples, and one 60 mg/kg/d sample. PCA showed an even clearer separation between dose levels than for miRNAs, with the vehicle and 50 mg/kg/d groups clustered somewhat closely and the 60 and 75 mg/kg/d groups displaying a gradient along principal component 1 (Fig. 5B). The first two principal components explained 69% of variance in the dataset (46% and 23%, respectively). The number of significant DE tRNA genes (Wald test, fold change ≥ 2, p-adj < 0.05) was 9, 117, and 204 in the 50, 60, and 75 mg/kg/d treatment groups, respectively, relative to control (Fig. 5C). After removing duplicate tRNA genes and comparing at the tRNA level, there were 2, 28, and 39 significant tRNAs at the 50, 60, and 75 mg/kg/d levels; at least one GluCTC and GlyCCC gene was significant at each of the three dose levels; and of the 24 additional tRNAs with at least one significant gene at the 60 and 75 mg/kg/d dose levels, GluTTC, LysCTT, AlaAGC, and CysGCA had the lowest adjusted p-values at 50 mg/kg/d (Fig. 5D). We generated coverage maps to analyze the tsRNA fragments present in the 2 consensus DE tRNAs and the next four most significant tRNAs (Fig. 5E), which showed the greatest coverage for GlyCCC, and mostly 5’ coverage for all except for GluTTC, which had more 3’ coverage. When analyzing coverage at the single-read level (Fig. 6), for GluCTC, LysCTT, AlaAGC, and Cys GCA, the vast majority of reads were 5’ fragments or halves, with a much smaller number of reads representing 3’ fragments. GluTTC reads consisted of an approximately 50:50 distribution of 5’ fragments and halves and 3’ fragments and halves. The predominance of the 5’ reads for most of tRNAs may be attributable to 5’ ligation bias resulting from modifications at 3’ ends of tRNAs (Upton et al. 2021; Scacchetti et al. 2024).

**Figure 5.**
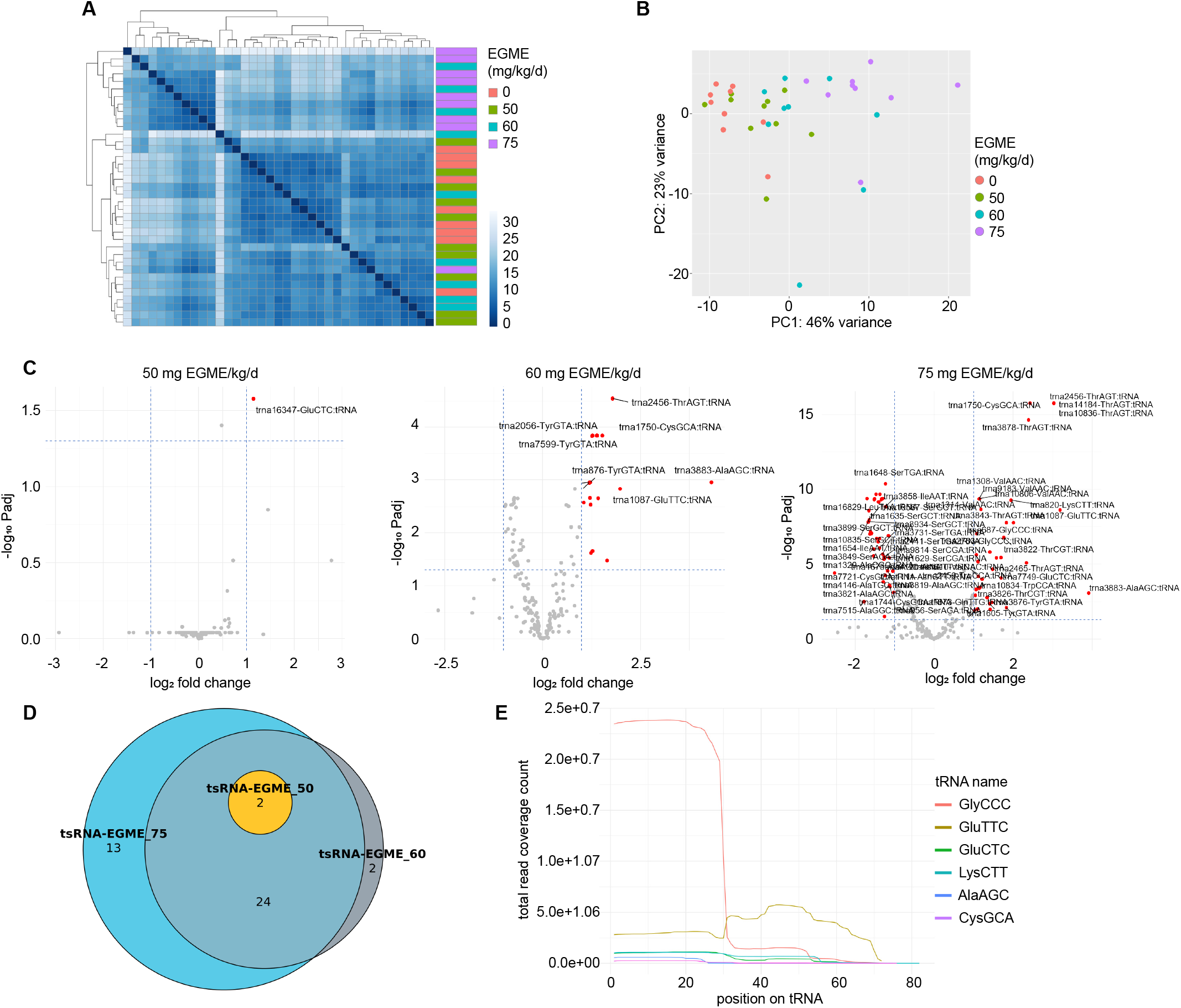
Changes in sperm tsRNA levels following EGME exposure. **A**. Hierarchical clustering of EGME- and vehicle control-treated samples showing Euclidean distance based on sperm tRNA read counts. **B**. Principal component analysis of EGME- and vehicle-control-treated samples based on sperm tRNA read counts. **C**. Differential expression of sperm tRNA between vehicle control and 50, 60, or 75 mg/kg/d EGME-treated samples (DESeq2 with the Wald test, p-adj < 0.05, fold change > 2). **D**. Venn diagram showing overlap of differentially expressed tRNAs in comparisons between control and 50, 60, and 75 mg/kg/d EGME, after removal of duplicate tRNA genes. Two unique tRNAs were common to all three treatment levels. **E**. Sequence coverage map showing read coverage of tRNA reads for the two tRNAs that were differentially expressed at all doses, GluCTC and GlyCCC, and the next four tRNAs with at least one significant gene at the 60 and 75 mg/kg/d dose levels, GluTTC, LysCTT, AlaAGC, and CysGCA, based on the lowest adjusted p-value at 50 mg/kg/d, scaled by read count on the y-axis.

**Figure 6.**
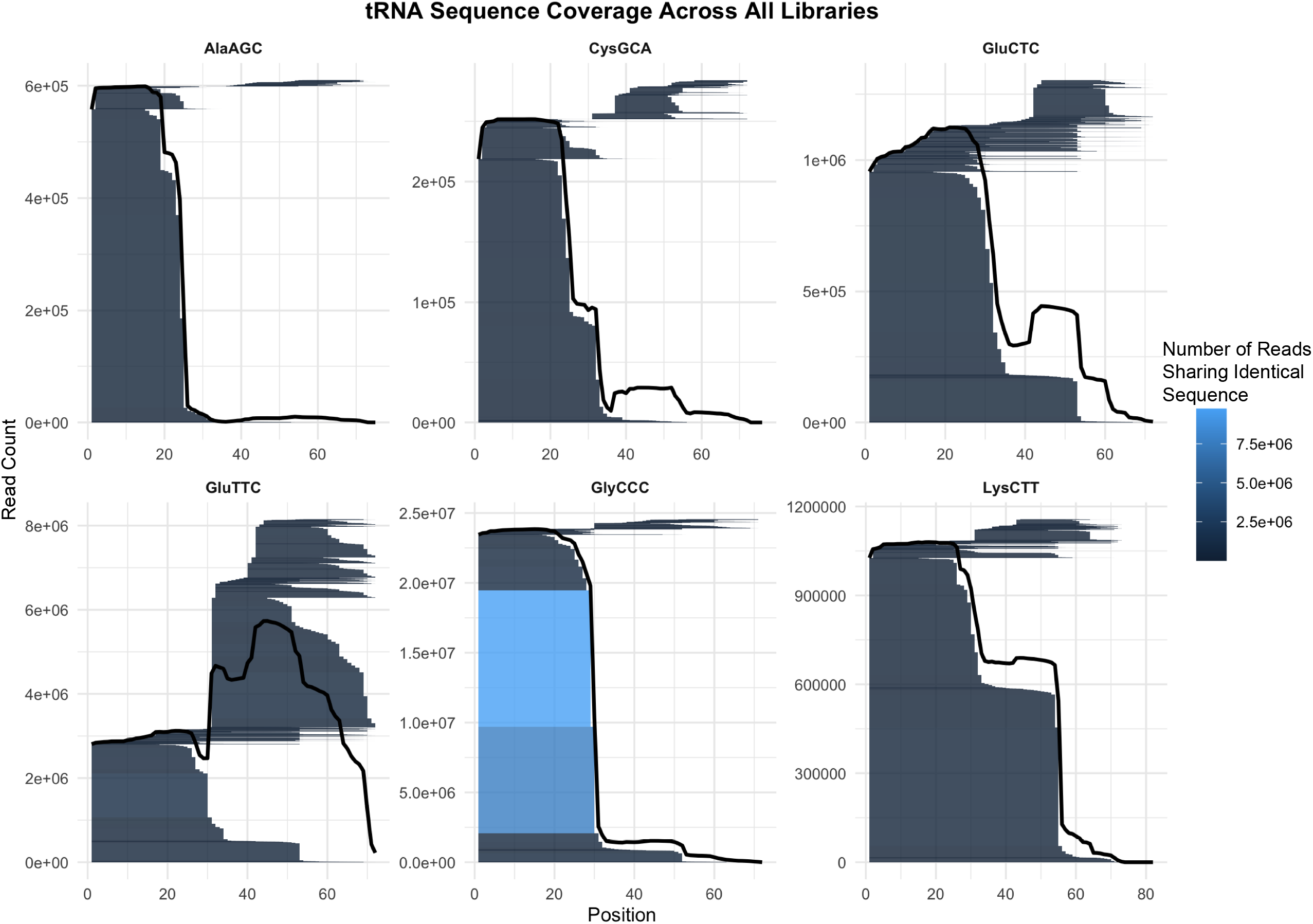
Detailed coverage maps for the most significant tsRNAs. Coverage maps showing all unique tRNA reads for the tRNAs that were differentially expressed at all doses, GluCTC and GlyCCC, and the next four tRNAs with at least one significant gene at the 60 and 75 mg/kg/d dose levels, GluTTC, LysCTT, AlaAGC, and CysGCA, based on the lowest adjusted p-value at 50 mg/kg/d. Plots show coverage of each individual read with color key to illustrate the number of copies of identical reads. The solid line shows the total read coverage at each site in the sequence. The majority of reads were 5’ fragments and halves, except for GluTTC, where more 3’ halves and fragments were present.

### EGME effects on sperm piRNAs

Small RNA reads mapped to 18,027 unique piRNAs in sRNAbench (Table S7), using annotations from piRBase (Wang et al. 2019). piRNAs separated by treatment group in unsupervised analyses, although less separation was visible in hierarchical clustering (Fig. 7A) than for other categories of RNAs. Similarly, PCA showed a gradient of values along PC1 with a cluster of 0, 50, and 60 mg/kg/d EGME samples mostly separating from a cluster of 75 mg/kg/d samples (Fig. 7B). The first two PCs explained 74% of the variance in the dataset (67% and 7% in the first two PCs, respectively). An increasing number of piRNAs were significant in DE analysis with each treatment group (Fig. 7C), with 1, 244, and 2,177 DE piRNAs in the 50, 60, and 75 mg/kg/d groups, respectively. A slight majority of the DE piRNAs were upregulated (148 out of 244 or 61%), relative to control, at 60 mg/kg/d, but at 75 mg/kg/d, 91.5% (1,991 of 2,177) of the DE piRNAs were downregulated relative to control. This large number of DE piRNAs is consistent with the size of this RNA category in spermatogenesis, which comprises more than 1 million piRNAs that are critical in the germline of rats and mice (Lau et al. 2006; Beyret et al. 2012; Watanabe et al. 2015) and typically diminishes as a fraction of the sncRNA reads with progression of spermatogenesis (Chen et al. 2016).

**Figure 7.**
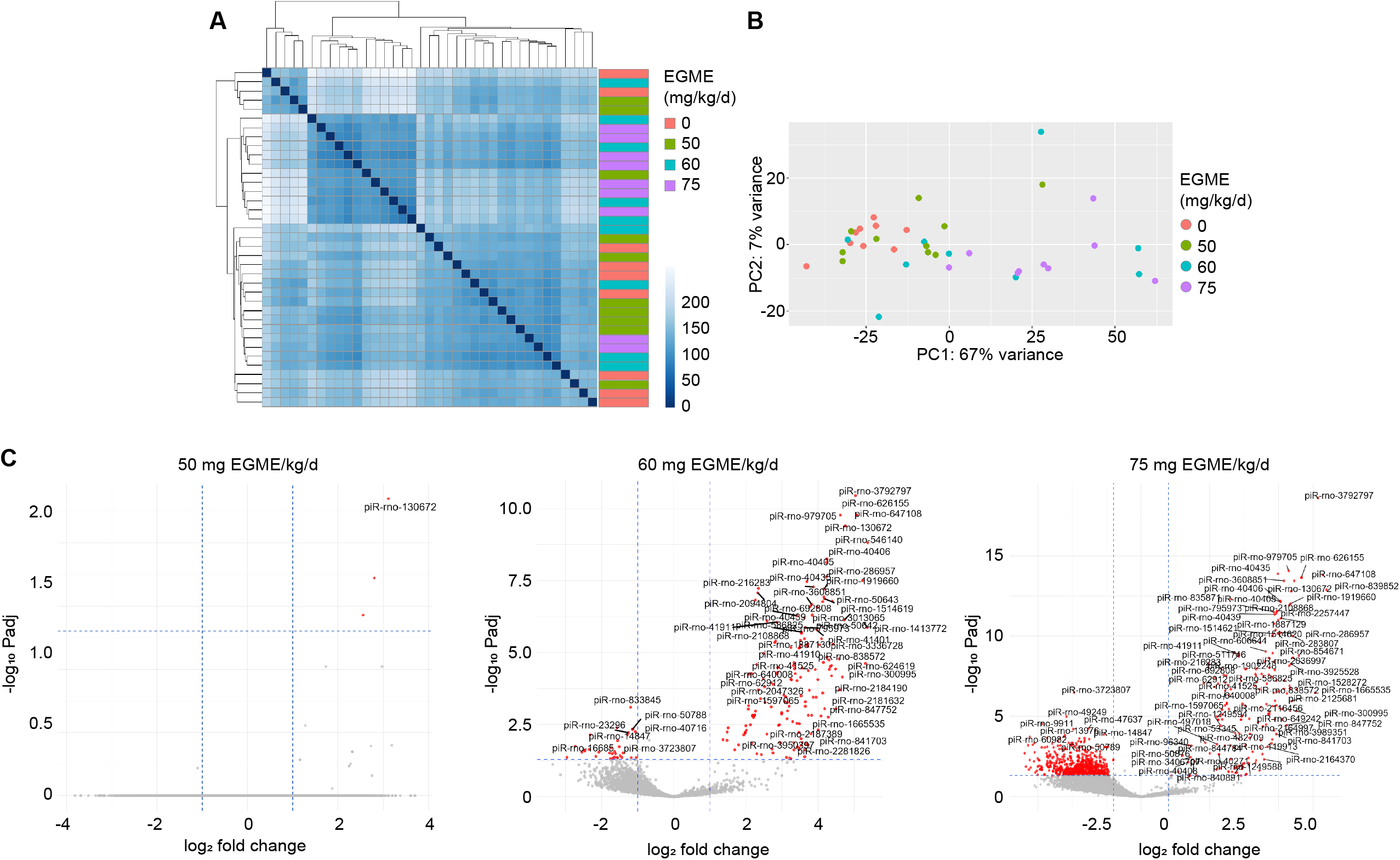
Changes in sperm piRNAs following EGME exposure. **A**. Hierarchical clustering of EGME- and vehicle control-treated samples showing Euclidean distance based on sperm piRNA read counts. **B**. Principal component analysis of EGME- and vehicle-control-treated samples based on sperm piRNA read counts. **C**. Differential expression of sperm piRNA between vehicle control and 50, 60, or 75 mg/kg/d EGME-treated samples (DESeq2 with the Wald test, p-adj < 0.05, fold change > 2).

### EGME effects on sperm rsRNAs

Small RNA reads mapped to the RNACentral rRNA annotations in sRNAbench were assigned to 8 unique rRNAs and analyzed for differential abundance using FUSION (Fig. 8, Table S8). In FUSION, all rsRNA reads are mapped to the parent rRNAs, which increases the power to detect differences, given that rRNAs are very large, and rsRNA reads for any one rRNA region may have low counts (Rawal et al. 2025).We found that 18S, 28S, 5S, 8S, LSU-S, and SSU-S rRNAs were significantly more abundant in EGME-treated rat sperm at all doses of EGME (p-adj < 0.05), while 12S and 16S rRNA reads were less abundant than vehicle control in the 60 and 75 mg/kg/d EGME treatment groups (Fig. 8A). These rsRNA reads did not appear to map randomly, with the exception of 28S-rRNA reads. 12S, 16S, and 18S rRNA reads mapped to a small number of overrepresented regions. 5S and 8S rRNA reads mapped predominantly to the 3’ half of the 5S rRNA sequence and the 5’ half of the 8S rRNA sequence (Fig. 8B).

**Figure 8.**
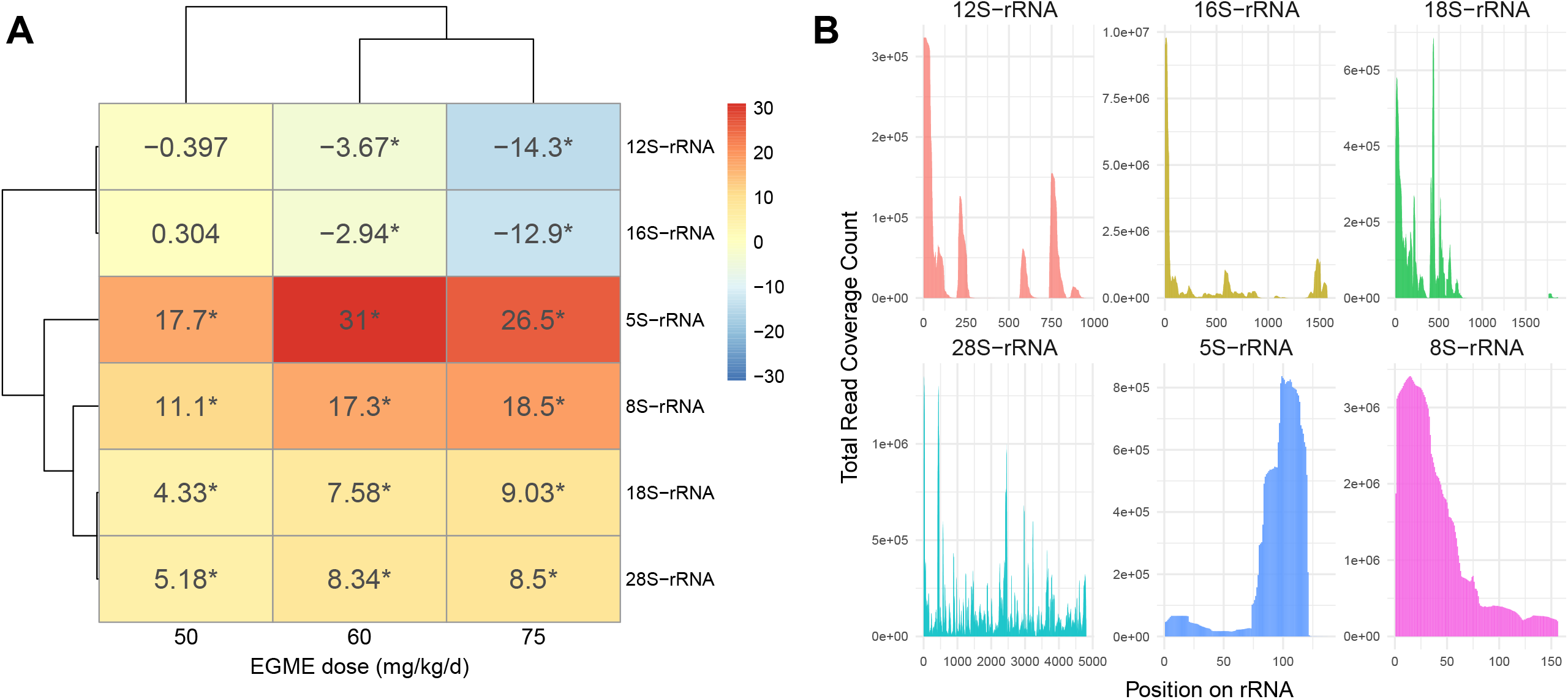
Changes in sperm rsRNAs following EGME exposure. **A**. Heatmap showing differential abundance of rsRNA reads in rat spermatozoa following EGME exposure. Heatmap color indicates the FUSION t-statistic from the comparison of rsRNA read counts for each EGME dose vs. vehicle control, with the t-statistic overlaid. *p-adj < 0.05. **B**. Coverage maps showing rsRNA read coverage for each EGME dose.

### EGME effects on other sperm small RNAs

EGME exposure level led to changes in sperm levels of multiple classes of small RNAs in addition to miRNAs, tsRNAs, and piRNAs (Table S9). These included rRNAs, mitochondrial rRNAs, snRNAs, lincRNAs, antisense RNAs, processed transcripts, and RNAs without known identities (identified as misc_RNA in Table S9). There was strong separation by unsupervised clustering analyses, similar to the pattern observed with miRNAs and tsRNAs. The hierarchical clustering result included two main clusters, one containing one 50 mg/kg/d sample, seven 60 mg/kg/d samples, and all nine 75 mg/kg/d samples; while the other cluster contained two 60 mg/kg/d samples, nine 50 mg/kg/d samples, and all nine vehicle control samples (Fig. S1A). PCA showed strong separation along the first PC. However, the first two PCs explained only 48% of the variance in the dataset (37% and 11%, respectively) (Fig. S1B). The number of DE RNAs in each group was 16, 79, and 133, in the 50, 60, and 75 mg/kg/d EGME groups, respectively (Table S9).

## Discussion

Ethylene glycol monomethyl ether (EGME), through its active metabolite, 2-methoxyacetic acid, is a testicular germ cell toxicant that causes testicular germ cell toxicity in the rat, with greater toxicity to spermatocytes than spermatogonia or spermatids (Chapin and Lamb 1984; Creasy and Foster 1984; Foster et al. 1984; Gray et al. 1985; Moss et al. 1985; Creasy et al. 1986), resulting in germ cell apoptosis (Ku et al. 1995). Meiotic spermatocytes are most sensitive to EGME, followed by pachytene spermatocytes, leptotene/zygotene spermatocytes, and step 1 round spermatids (Creasy and Foster 1984; Foster et al. 1984).The no observed adverse effect level (NOAEL) dose for germ cell death has been reported as 50 mg/kg/d following 11-day exposure to EGME and 250 mg/kg EGME in a single-dose experiment (Creasy et al. 1986). EGME-induced germ cell death is preceded by mitochondrial damage 16 hours after a single 500 mg/kg dose (Foster et al. 1983).

Because EGME selectively targets spermatocytes, an exposure duration of as little as five days results in decreased male fertility that becomes apparent three to four weeks after cessation of exposure, with the greatest magnitude of effect occurring in weeks five to six post-exposure, with a LOAEL of 100 mg/kg/d (Chapin, Dutton, Ross, and Lamb 1985; Chapin, Dutton, Ross, Swaisgood, et al. 1985; Holloway et al. 1990). The delayed fertility effect corresponds with the 4-5 weeks required for spermatocytes to mature into sperm. A dose of 200 mg/kg/d has also been reported to increase the number of resorptions five to six weeks after a five-day exposure (Chapin, Dutton, Ross, and Lamb 1985), suggesting that paternal EGME exposure can cause developmental toxicity without exposure of the dam or embryo. There is extensive evidence that EGME is non-genotoxic. EGME is negative in the Ames assay, with and without metabolic activation by S9 fraction (Hoflack et al. 1995). It is also negative in a variety of other screens including the unscheduled DNA synthesis test in human fibroblasts and the bone marrow metaphase analysis in rats (McGregor 1984). However, EGME has been described as potentially genotoxic because of positive results in an assay called the dominant lethal test in rats (McGregor et al. 1983; McGregor 1984). The dominant negative test is a screen for pre- and post-implantation embryonic loss caused by exposure of a parent, in this case the father, to a chemical. The theory of the test is that DNA damage will lead to dominant lethal mutations, where a mutant copy derived from only one parent will result in early embryonic loss (Bateman 1973). Notably, the theory of this test rests on the concept that mutations do not cause cytotoxicity until the next mitotic division, which means that stable DNA damage in gametes can cause embryonic failure, even if it did not prevent fertilization. Therefore, the belief is that dominant lethal mutations are detected when the test article damages post-meiotic germ cells, which in the testis means spermatids or spermatozoa (Bateman 1973). EGME, conversely, causes damage preferentially to the spermatocyte, which has been observed histologically (Creasy and Foster 1984) and in ultrastructure analysis by electron microscopy (Creasy et al. 1986; Anderson et al. 1987; Lee and Kinney 1989). Therefore, EGME is likely to cause post-implantation embryonic loss through some mechanism other than dominant lethal mutations. Given our results, one possibility is that this occurs through dysregulation of sncRNA-mediated control of embryonic transcription or translation. Alternatively, if paternal EGME exposure does not directly cause DNA damage, it is possible that it does introduce dominant lethal mutations through a mechanism other than direct germ cell DNA damage, such as loss of piRNA-mediated control of genomic stability. This possibility cannot be excluded completely: although EGME does not directly induce DNA strand breaks, to our knowledge, there are no published tests of sperm DNA damage in EGME-treated animals. EGME does alter sperm parameters. For example, in the experiment by Stermer et al. (2019), 75 mg EGME/kg/d caused a decrease in sperm motility. An inhalation dose of 500 ppm EGME, but not 25 ppm EGME, for 7 hours/day for a period of 5 days induced sperm abnormalities, especially amorphous head defects (McGregor et al. 1983). Similarly, oral exposure to 100 or 200 mg/kg/d EGME caused significant increases in abnormal sperm, mostly consisting of head defects (Chapin, Dutton, Ross, and Lamb 1985). These defects may be consistent with impaired sperm maturation resulting from EGME exposure. In our dataset, downregulation of the 12S and 16S mitochondrial rRNA sequences (Fig. 8) may also be consistent with an overall suppression of sperm maturation by EGME, possibly affecting mitochondrial content or translation capacity. However, these hypotheses require further investigation and experimental validation before any conclusions can be drawn.

Stermer et al. (2019) used low-dose EGME as a model to understand how sub-cytotoxic germ cell effects of EGME could alter markers of reproductive health. Exposure of male rats to 50-75 mg/kg/d EGME for five days, followed by a five-week withdrawal, resulted in reduced sperm count and increased retained spermatid head (RSH) count, with a NOAEL of 60 mg/kg/d and a LOAEL of 75 mg/kg/d (Stermer et al. 2019). In addition to these apical effects, all doses of EGME that were tested (50-75 mg/kg/d) altered the proportion of miRNA, piRNA, and tsRNA reads in sperm and the distribution of small RNA read lengths. An ROC curve analysis revealed that average sperm small RNA read length was a better EGME dose classifier than sperm motility or RSH. In the present paper, we performed a new analysis of these data to identify the sperm small RNAs that were differentially abundant following EGME exposure, to obtain novel mechanistic insights about EGME-driven changes in spermatogenesis and to identify RNAs that may contribute to differences in embryonic development following paternal EGME exposure. We identified EGME dose-dependent changes in abundance of sperm miRNAs (Fig. 1-2), tsRNAs (Fig. 5-6), piRNAs (Fig. 7), and other classes of small RNA (Fig. 8).

miRNAs canonically act as negative translational regulators of their mRNA targets, but they can act as positive regulators in some instances (O’Brien et al. 2018). There is evidence that sperm miRNAs regulate pre-implantation embryonic transcription in the mouse, at the 4-cell and blastocyst stages (Conine et al. 2018). In our analysis, the number of significant differentially expressed (DE) miRNAs increased with EGME dose (Fig. 1), and there were 12 DE miRNAs present in all three doses, which all showed a monotonic increase in abundance with dose, and which were highly reproducible based on a Monte Carlo analysis (Fig. 2). These 12 miRNAs include members of the Mir-23-27-24 cluster and the Mir-200 family. The Mir-200 family consists of five members, including Mir-200a, 200b, and 429, which were DE in our study and have roles in epithelial development during embryogenesis (Korpal and Kang 2008; Sundararajan et al. 2022). The Mir-23-27-24 cluster, members of which were also upregulated by EGME in this experiment, is known to regulate stem cell differentiation during embryonic development (Yap and Chen 2024). Consistent with the known roles of these miRNAs in development, we identified over 1,000 high confidence targets of the 12 core EGME dose-dependent miRNAs, many of which are involved in developmentally important biological processes and expressed in human preimplantation embryos (Fig. 3B-D, Table S2-S3). Several of the EGME dose-dependent sperm miRNAs have known functions in spermatogenesis, and/or embryo development, including miR-221-3p, miR-27a-3p, miR23a-3p, and let-7b-5. miR-221-3p regulates germ cells to maintain undifferentiated spermatogonia states (Yang et al. 2013). miR-27a-3p is overexpressed in azoospermic men (Norioun et al. 2020), and miR-23a/b-3p are overexpressed in subfertile men (Abu-Halima et al. 2019). Let-7 miRNAs regulate germ cell transcription, with let-7b being especially highly expressed in pachytene spermatocytes (Sangiao-Alvarellos et al. 2015). We hypothesize, therefore, that levels of miRNAs such as let-7b could be altered in sperm following an injury to spermatocytes and could influence embryo development as a result. In addition to those miRNAs that have previously characterized roles in male reproductive biology, it is notable that let-7c-5p, miR-200a-3p, and miR-200b-3p increased with EGME dose in our dataset (Fig. 2), but decreased during epididymal maturation in the mouse in Sharma et al. (2018).

These changes in sperm miRNA levels are consistent with a previous report that 300 mg/kg/d EGME exposure for 4 days in the cynomolgus monkey led to downregulation of 16 miRNAs and upregulation of 347 miRNAs in the testis (Sakurai et al. 2015). Although it is difficult to compare directly across research models and compare a testis dataset with a sperm dataset, one major similarity is that the preponderance of DE miRNAs in both the Sakurai study and our study (Fig. 2C) were upregulated. The authors of the Sakurai study reported a list of the 18 upregulated and 7 downregulated miRNAs with the largest fold difference, but there were few matches with our results. rno-miR-365-3p and hsa-miR-365a-5p were upregulated in the respective datasets. Similarly, miR-34b-5p and miR-34c-3p were downregulated in the Sakurai et al. study, and in our dataset, miR-34b-3p and miR-34c-3p were downregulated, while miR-34a-5p was upregulated only in the 60 mg/kg/d group (Table S1). The lack of direct overlap between the two datasets is perhaps unsurprising because the monkey miRNAs were annotated using human data from miRbase, which includes 1,917 human miRNAs and only 496 rat miRNAs. Inherent species differences or differences in current knowledge of miRNA genes and active miRNAs between species may contribute to this RNA-level difference, although the overall trend was similar between the two studies.

Although miRNAs are well-characterized, recent evidence shows that the majority of the RNAs in sperm are tsRNAs and rsRNAs, which are underrepresented in sequencing results because of modifications that impede reverse transcription (Shi et al. 2021; Hernandez et al. 2023). Although the sequencing approach from which the present EGME dataset is derived did not use the PANDORA-seq methodology, a significant proportion of the reads in this dataset mapped to tRNA sequences: 51.4-65.5%, depending on EGME dose (Stermer et al. 2019). Consistent with that finding, we identified 41 DE tRNAs, including 2 that were DE in all EGME doses, GluCTC and GlyCCC (Fig. 5). Coverage maps showed that the tsRNAs that contributed to these differential read counts were predominantly 5’ tRNA fragments and halves (Figs. 5-6). Notably, there is a known bias toward 5’ tRNA halves in library preparation (Upton et al. 2021), which may account to some extent for this coverage pattern. However, some 3’ halves and fragments were detected for all tRNAs, especially for GluTTC (Fig. 6). This is consistent with previous observations on human sperm using similar sequencing methodology (Mehta and Singh 2025).

tsRNAs may have multiple functions, including miRNA-like RNAi functions, as well as aptamer-like functions that are dictated by their three-dimensional structures, such as RNA-protein binding that dictates RNA binding protein localization, can stabilize transcripts, and can induce stress granule formation (Chen and Zhou 2023; Kuhle et al. 2023). tsRNAs may also participate in regulation of transposable elements (Sharma et al. 2016). A particularly important role in the context of our current study is repression of pyrimidine-enriched sequence translation in embryonic stem cells (Guzzi et al. 2018; Kuhle et al. 2023). Although the understanding of tsRNA functions is still emerging, they likely perform several important regulatory functions. With respect to our dataset, recent evidence shows that the sperm quantity of one of our DE tsRNAs, 5’tRF-GluCTC, is negatively associated with ART outcomes in humans (Hekim et al. 2025).

Similarly, many reads in our dataset mapped to rsRNA sequences (Fig. 8). Interest in rsRNAs has expanded in recent years, with the discovery that they were underrepresented in many small RNA sequencing datasets. The rsRNA population in sperm is influenced by *Dnmt2*, which suggests a role for rsRNAs in epigenetic programming (Zhang et al. 2018). rsRNAs in combination with tsRNAs regulate the transcriptome of mouse embryonic stem cells (Shi et al. 2026), which is evidence that both classes have the potential to regulate transcription during early embryonic development. Moreover, sperm rsRNAs are responsive to factors that influence male reproduction. The population of rsRNAs in mouse and human sperm changes with increasing age (Shi et al. 2026). Further, human sperm rsRNAs are upregulated after 1 week of high-sugar diet (Nätt et al. 2019), and exposure to a low dose of dicyclohexyl phthalate (10 mg/kg) for four weeks increased mouse sperm rsRNA counts (Liu et al. 2023). This suggests that rsRNAs may be an important category of sperm RNAs that could mediate the effects of paternal exposure on offspring.

EGME caused a massive increase in the number of upregulated piRNAs from 1 significant DE piRNA at 50 mg/kg/d to over 1000 at 75 mg/kg/d (Fig. 7). piRNAs are important for maintenance of germline genomic integrity by regulating insertion of transposable elements (Wang and Lin 2021). piRNAs also control some aspects of posttranscriptional processing and cleavage of RNAs (Watanabe and Lin 2014) and have a variety of other epigenetic control mechanisms (Ku and Lin 2014). Hypothetically, differential expression of piRNAs could indirectly destabilize the genome of germline cells, despite EGME not being a direct mutagen (Hoflack et al. 1995). This hypothesis would require further testing, but it presents a possible sperm sncRNA-mediated mechanism of paternal EGME effects on embryonic development. As noted above, rsRNAs are also prevalent in sperm, representing over half of sperm RNA reads when PANDORA-seq is employed (Shi et al. 2021; Chen and Zhou 2023). We also analyzed other sperm sncRNAs, such as rRNAs, mtRNA, snRNA, and lincRNA, and we found that the number of DE RNAs increased across those classes with increasing EGME dose (Fig. S1).

Overall, we found that short-term exposure to low doses of EGME led to changes in sperm RNA that were detectable 5 weeks after cessation of exposure, consistent with the known progression of EGME-induced testicular injury and reproductive performance in rats (Chapin, Dutton, Ross, and Lamb 1985). This is also consistent with prior reports that EGME exposure altered testicular gene and protein expression (Wang & Chapin, 2000, Fukushima et al., 2005, Yamamoto et al., 2005) and miRNA expression in rats and non-human primates (Fukushima et al., 2011, Sakurai et al., 2015). EGME’s property of targeting pachytene spermatocytes may be the reason for its potent and persistent effects on testicular germ cell RNA expression, as the peak of RNA synthesis during spermatogenesis occurs in the pachytene spermatocytes (Monesi, 1965). However, testicular toxicants with a variety of target cells and toxicity mechanisms have been reported to alter sperm mRNA expression. These include pharmaceuticals (Dere et al. 2017) and industrial or environmental testicular toxicants including the germ cell toxicant, carbendazim, and the Sertoli cell toxicant, 2,5-hexanedione (Pacheco et al. 2012). There is also a recent report that phthalate exposure altered miRNA, tsRNA, and piRNA levels in epididymal extracellular vesicles (Oluwayiose et al. 2025), some of which are transferred to sperm (Sharma et al. 2018). Similarly, a 12-week exposure to a mixture of per- and polyfluoro alkyl substances (PFAS) resulted in increased numbers of sperm tsRNAs, miRNAs, and piRNAs at doses that caused few phenotypic changes in adults and no changes in fertilization rate (Gillespie et al. 2025). Other parental treatments alter sperm miRNAs with apparent intergenerational biological effects, including maternal stress (Gapp et al. 2014), paternal exercise (Yin et al. 2025 Oct), and paternal metabolic disorders (Zhang et al. 2018; Zhang et al. 2019).

In humans, there is evidence that sperm RNAs may act as markers of fertility or exposures that affect fertility. For example, in one study the total amount of RNA per sperm cell differed based on alcohol use (Bianchi et al. 2019). We recently reported that human sperm mRNA profiles differed between study participants with different clinical fertility status (Qi et al. 2025). Another recent study found that miRNAs, including let-7b-5p, differed between proven fertile and subfertile men (Abu-Halima et al. 2024). Here, our new analysis identified changes in rat sperm sncRNA following EGME exposure, which is consistent with the hypothesis that paternal exposure to environmental toxicants can alter sperm RNA profiles and that this is a putative mechanism of reduced male fertility. These changes affected RNAs known to have roles in human fertility. Almost all of the 12 DE miRNAs were consistently present in normozoospermic fertile human sperm across two major studies (Hua et al. 2019; Abu-Halima et al. 2024), while some of these were also reported in normozoospermic fertile men from India (Mehta and Singh 2025), USA (Morgan et al. 2020), and Spain (Pantano et al. 2015), suggesting their critical significance to human fertility. Future experiments should test the role of specific EGME dose-dependent candidate miRNAs on rat embryo development and other sncRNA-mediated epigenetic toxicity mechanisms. In summary, EGME not only sensitively alters the proportion of different classes of small RNAs present in sperm, as previously reported (Stermer et al. 2019); it also leads to dose-dependent changes in specific small regulatory RNAs in sperm. Investigation of the roles of those sncRNAs in spermatogenesis and embryo development may lead to discovery of novel EGME toxicity mechanisms.

## Data Availability Statement

Stored in repository: The data that support the findings of this study are openly available in NCBI databases. The RNA sequence reads analyzed in this study are available form the NIH Sequence Read Archive database, accession no. PRJNA492909. The raw and normalized data supporting the RNA-seq analysis are available in the NCBI Gene Ontology Omnibus (GEO) database, accession number GSE313131. The analysis code is publicly available at https://github.com/compbiocore/egme.

