## Supplementary material for "Ethylene Glycol Monomethyl Ether Altered Rat Sperm Small RNAs with Critical Developmental Roles": Fig. S1

**Contents of this document include supplemental figure S1.**

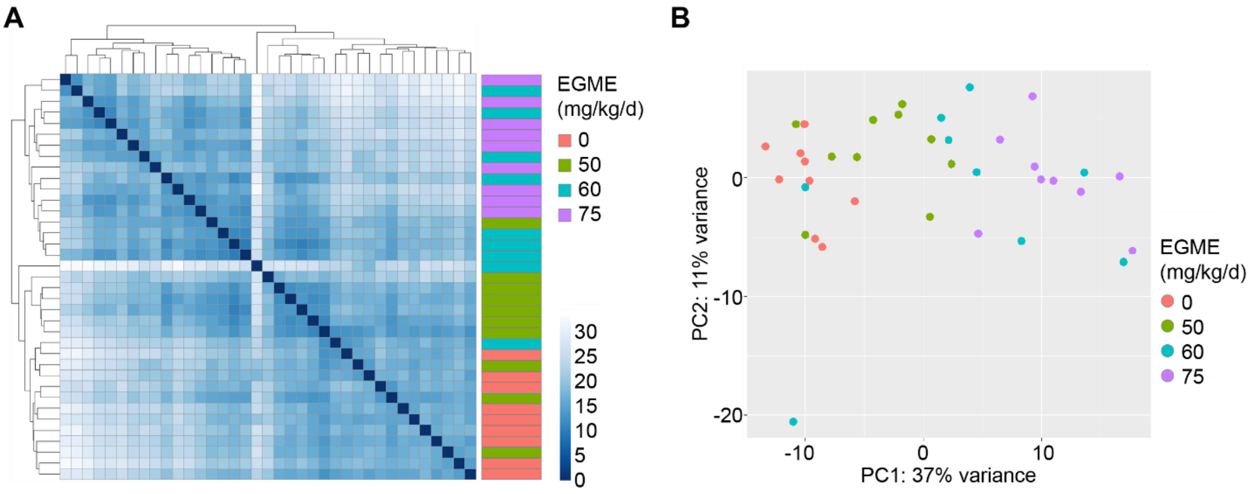


**Figure S1. Changes in other sperm small RNAs following EGME exposure.**

**A.** Hierarchical clustering of EGME- and vehicle control-treated samples showing Euclidean distance based on remaining sperm read counts. After separating miRNA, tRNA, and piRNA reads, the remaining DE RNAs were from RNA classes including rRNA, mitochondrial rRNA, lincRNA, snRNA, **B.** Principal component analysis of EGME- and vehicle-control-treated samples based on sperm sncRNA read counts.
